# BrightEyes-Scope: A Reference Implementation for Image Scanning Microscopy with SPAD Array Detection

**DOI:** 10.64898/2026.09.30.755689

**Authors:** Laurent Le, Alessandro Zunino, Eli Slenders, Andrea Bucci, Mattia Donato, Giuseppe Vicidomini

**Author notes:** **Correspondence** Giuseppe Vicidomini. these authors contributed equally (Laurent Le, Alessandro Zunino). **Funding information** This work was funded by PNRR MUR-M4C2–Action 1.4—Call Potenziamento strutture di ricerca e creazione di ‘campioni nazionali’ di R&S (CUP J33C22001130001), National Center for Gene Therapy and Drugs based on RNA Technology no. CN00000041, and by the European Research Council, ERC-StG grant No. 101219234 – MotionPicture (ES) and ERC-CoG No. 818699 – BrightEyes (GV). Views and opinions expressed are however those of the author only and do not necessarily reflect those of the European Union or the European Research Council Executive Agency. Neither the European Union nor the granting authority can be held responsible for them.

## Abstract

Image scanning microscopy (ISM) is emerging as the natural evolution of confocal laser-scanning microscopy, combining optical sectioning and a two-fold improvement in lateral resolution with high signal-to-noise ratio. Single-photon avalanche diode (SPAD) array detectors enable an effective implementation of ISM by recording a micro-image at each scan position with single-photon sensitivity, negligible read-out noise, and without the fixed frame-readout constraints of conventional cameras. Nevertheless, SPAD array-based ISM remains largely confined to laboratories with expertise in microscope development. Here, we present BrightEyes-Scope, a reference implementation of a SPAD array-based ISM system. We provide step-by-step guidelines for optical design, system construction, alignment, calibration, data acquisition, and image reconstruction. Presented as a complete ISM microscope, BrightEyes-Scope also serves as a reference for upgrading existing laser-scanning microscopes. We anticipate that it will accelerate the adoption of ISM for quantitative super-resolution imaging.

## 1. INTRODUCTION

Fluorescence microscopy is a central tool in biological imaging thanks to its molecular specificity and sensitivity [1]. Two imaging architectures dominate modern fluorescence microscopy: widefield microscopy (WFM), in which the entire field of view is excited simultaneously and the emitted fluorescence is recorded by a camera detector, and laser-scanning microscopy (LSM), in which the sample is sequentially excited by a diffraction-limited focused beam and the emitted fluorescence is detected by a single-point detector. In both cases, the achievable spatial resolution is fundamentally limited by diffraction. In WFM, image formation is governed by the emission point spread function (PSF), whereas in LSM it is primarily determined by the excitation PSF. Additionally, neither conventional WFM nor conventional LSM provides optical sectioning. In the WFM architecture, structured illumination microscopy (SIM)first introduced optical sectioning through patterned illumination [2] and was later extended to achieve a two-fold improvement in lateral resolution beyond the diffraction limit by appropriately tailoring the illumination pattern [3]. In LSM architecture, the analogous development is confocal laser-scanning microscopy (CLSM), which introduces a pinhole in the detection path to reject out-of-focus fluorescence, thereby providing optical sectioning [4, 5]. In principle, reducing the pinhole diameter also narrows the effective confocal PSF, allowing up to a two-fold improvement in lateral resolution with respect to the diffraction limit, analogous to SIM. However, this theoretical resolution gain is achieved only with an almost infinitesimally small pinhole, which drastically reduces the detected photon flux and consequently the signal-to-noise ratio (SNR). As a result, conventional confocal microscopy provides optical sectioning but is not generally operated as a practical super-resolution technique.

Image scanning microscopy (ISM) overcomes this fundamental compromise by replacing the single-point detector of a confocal microscope with a detector array [6, 7]. Each detector element is much smaller than one Airy unit (A.U.) and records an image equivalent to that acquired with an almost-closed pinhole. At the same time, the detector elements collectively span approximately one Airy unit, ensuring efficient photon collection and therefore a high signal-to-noise ratio (SNR), while preserving the optical sectioning capability. Computational reconstruction algorithms [8] then combine the individual detector images into a single image with enhanced lateral resolution. Moreover, recent reconstruction methods have demonstrated that optical sectioning can be preserved even when the detector array extends beyond one Airy unit, further improving the SNR while maintaining the sectioning capability of ISM [9, 10, 11]. Together with the development of fast detector arrays,first exemplified by Airyscan detectors [12] and more recently by asynchronous read-out single-photon avalanche diode (SPAD) arrays [7], these advances have established ISM as a practical imaging modality capable of simultaneously providing optical sectioning, high photon efficiency, and an approximately two-fold improvement in lateral resolution.

Today, ISM is emerging as the natural evolution of confocal laser-scanning microscopy and is progressively being integrated into commercial laser-scanning microscope platforms, including Nikon NSPARC, Zeiss Airyscan, Abberior Mirava Polyscope, PicoQuant Luminosa, and Genoa Instruments PRISM. Because ISM preserves the excitation, scanning, and imaging architecture of a conventional confocal microscope, it remains fully compatible with the fluorescence labelling strategies, sample preparation protocols, and experimental workflows routinely employed in CLSM. This architectural continuity also makes its implementation remarkably straightforward, as the only fundamental hardware modification is the replacement of the conventional single-point detector with a detector array. Consequently, many existing laser-scanning microscopes, particularly custom-built systems, can be readily upgraded to ISM while preserving most of their original optical architecture.

For this reason, we present a detailed and practical tutorial describing the implementation of a simple – yet powerful – ISM system. The tutorial provides a step-by-step guide to the design, construction, alignment, calibration, data acquisition, and image reconstruction of a complete ISM microscope. Although the implementation is presented as a microscope built from scratch, it is equally intended to serve as a reference for understanding the design principles, hardware modifications, and practical challenges involved in converting an existing laser-scanning microscope into an ISM system. The implementation presented here employs a 100× NA 1.4 oil-immersion objective, 488 nm excitation, and fluorescence detection in the green spectral range, the same methodology is readily applicable to a broad range of microscope configurations. While ISM can be implemented using different types of pixelated detectors [6, 12, 7], we focus on a SPAD array detector, which has become the detector of choice owing to its single-photon sensitivity, without readout noise [13], and its synergy with important advanced fluorescence microscopy methods typically combined with confocal laser-scanning microscopy, such as fluorescence lifetime imaging [14, 10, 11, 15] and fluorescence correlation spectroscopy [16, 17].

## 2. MICROSCOPE ASSEMBLY

Image-scanning microscopy shares the excitation and beam-scanning architecture of a CLSM. Its principal distinguishing feature lies in the fluorescence-detection path, where the pinhole and single-element detector are replaced by an array that records a micro-image of the illuminated region at each scan position. Accordingly, its implementation is governed by three main optical requirements. First, the galvanometric scanners must be conjugated to the objective back focal plane. In this configuration, rotation of the galvanometric mirrors changes the illumination angle at the objective without translating the beam across its back focal plane. This enables telecentric scanning and minimizes scan-dependent variations in illumination intensity, focal position, and optical aberrations. Second, analogously to the positioning and sizing of the pinhole in a confocal laser-scanning microscope, the sample plane must be imaged onto the detector array with an appropriate magnification. This magnification should be selected such that each detector element acts as a sub-Airy-unit pinhole, whereas the complete array spans a sufficiently large detection area to collect most of the emitted fluorescence. Third, the excitation focus and the detection PSFs associated with the individual detector elements must be correctly registered in the sample plane. Together, these conditions ensure that the detector elements record the spatially shifted scanned images required to reconstruct a super-resolved image with a high signal-to-noise ratio. The construction sequence described below is designed to establish these conditions progressively while keeping each alignment step independently verifiable. We first define the objective position and construct the telecentric scanning relay; we then introduce and align the excitation beam;finally, we assemble the detection path and position the SPAD array.

The system is built on a 60 × 45 cm breadboard to provide a minimal, complete, and compact implementation that can also be transported for workshops and practical demonstrations. Although the system is intentionally limited to a single excitation wavelength and one detection channel, the same design and alignment principles can be readily extended to more advanced configurations incorporating multiple lasers and spectral detection channels. A cage-based construction is used to improve mechanical stability and facilitate alignment. The cage architecture constrains the optical elements to a common axis, simplifies the implementation of orthogonal beam paths, and allows components to be translated along the optical axis while approximately preserving their lateral centering.

### 2.1. Aligning the scanning module

In our implementation, construction begins with the objective lens (OL; 100×/1.4 NA oil Plan Apo, Leica), which is mounted in an inverted-microscope configuration. The objective defines the sample plane, optical axis, and geometrical constraints of the microscope; its position is therefore used as the fixed reference for assembling the remaining optical components. The sample is mounted on a three-axis flexure stage (Thorlabs MAX302/M NanoMax Stage with 3 Differential Adjuster) that provides both coarse mechanical positioning and fine piezoelectric control along X, Y, and Z. Consequently, the dimensions, travel range, and accessibility of the complete sample-positioning assembly must be considered before fixing the objective position and constructing the surrounding optical relay.

Two steering mirrors (M1; Thorlabs KCB1/M and M2; Thorlabs KCB2EC/M) are placed between the objective and the tube lens (TL; Leica, f = 200 mm). By beam walking, these mirrors provide independent control over the beam position and propagation angle, allowing the excitation beam to be centered on the objective back aperture and incident perpendicular to it when the galvanometric mirrors are in their zero-offset position. Their locations should therefore be selected to permit convenient adjustment without interfering with the sample stage or other microscope components.

The objective and tube lens form the first optical relay and must be arranged as an afocal pair. The tube lens must therefore be compatible with the infinity-corrected objective and positioned so that the back focal plane of the objective is conjugated to the front focal plane of the tube lens. Incorrect spacing between these components can introduce defocus and field-dependent aberrations, which subsequently affect both scanning telecentricity and image quality. To establish this conjugation, we use a portable alignment laser (Thorlabs PL201), which can be introduced at different positions in the cage system, together with a shearing interferometer (Thorlabs SI254). The shear plate should be selected according to the beam diameter and positioned where the beam is sufficiently expanded to produce clearly distinguishable interference fringes. For the objective–tube-lens alignment, the collimated alignment beam is injected through the objective, and the shearing interferometer is placed after the tube lens, where the beam diameter is increased by the angular-magnification factor of the afocal pair. The objective is kept fixed, whereas the tube lens is translated along the optical axis until the interferometer indicates that the emerging beam is collimated (Figure 2a). This procedure establishes the correct axial spacing of the objective and tube lens; lateral centring is verified separately by ensuring that the beam propagates along the mechanical axis of the cage system.

**FIGURE 1.**
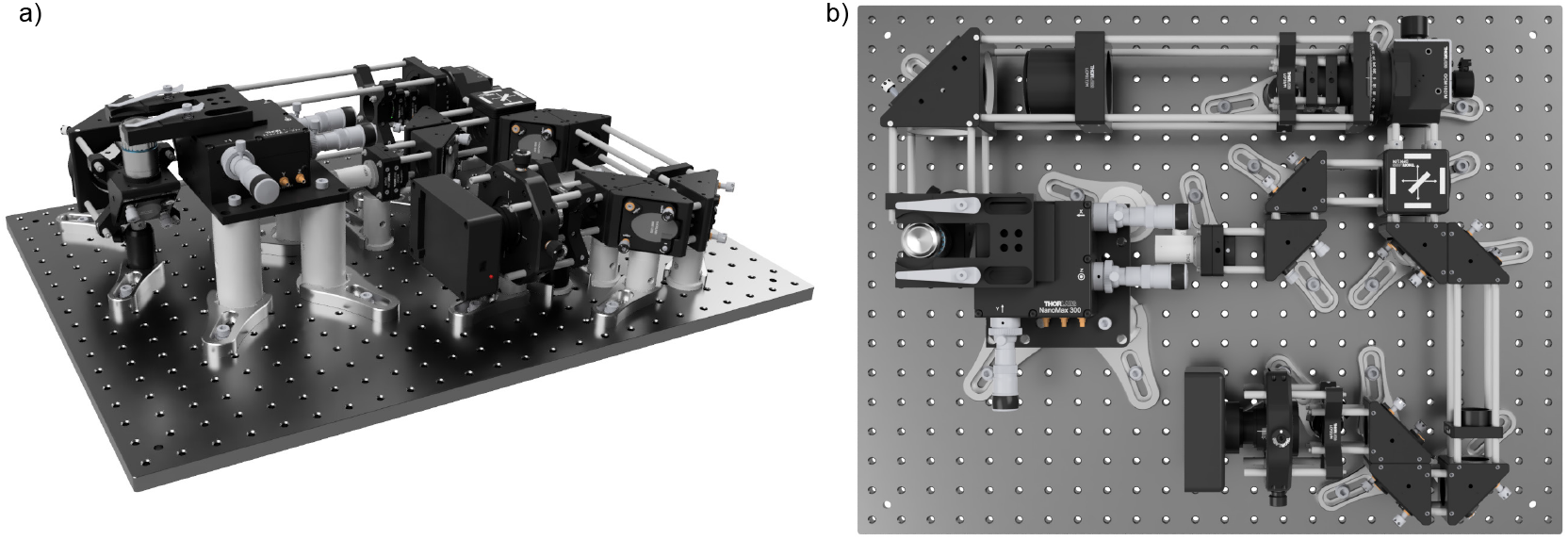
Optical configuration of BrightEyes-Scope. (a) Three-dimensional representation of the microscope assembly. (b) Top view of the microscope. The spacing between M6 holes is 25 mm.

**FIGURE 2.**
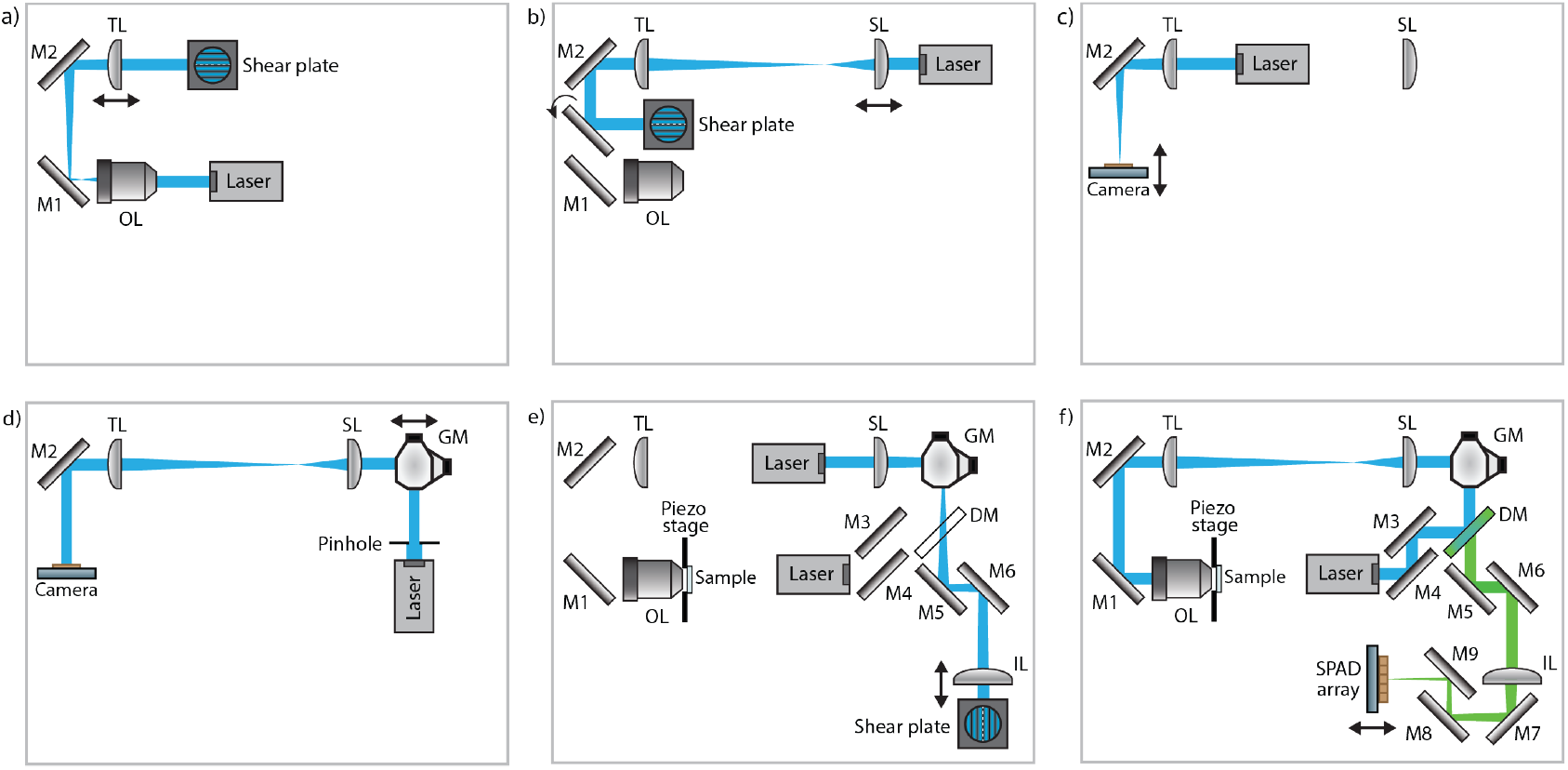
Sequential alignment of the microscope optical system. Schematic representation of the principal steps used to establish the required conjugate planes and align the excitation and detection paths. OL: objective lens; M1–M9: mirrors; TL: tube lens; SL: scan lens; GM: galvanometric mirrors; DM: dichroic mirror; IL: imaging lens.

The scan lens (SL; f = 50 mm, Leica SP5) is then positioned relative to the tube lens. This scan lens was used to acquire all the data presented in this manuscript; however, we also successfully tested a commercially available scan lens with the same focal length (Thorlabs CLS-SL). Together, the scan lens and tube lens relay the angular deflections produced by the galvanometric mirrors to the objective back focal plane, thereby enabling approximately telecentric scanning at the sample. A scan lens designed to minimize field-dependent aberrations and scan distortion is therefore preferable. An F-theta design can additionally improve the linearity between the galvanometer deflection angle and the displacement of the excitation focus, particularly over a large field of view, but does not by itself guarantee telecentricity. The scan lens is translated along the optical axis until the scan-lens–tube-lens pair forms an afocal system, as verified using the shearing interferometer (Figure 2b).

The galvanometric scanning unit (GM; Thorlabs GVS002) is subsequently integrated into the optical relay. Ideally, the pivot points of both galvanometric mirrors would lie in the front focal plane of the scan lens. Their angular deflections would then be mapped onto angular deflections at the objective back focal plane without producing a scan-dependent translation of the beam across the objective aperture. In a two-axis galvanometric scanner, however, the mirrors are necessarily separated along the optical path and cannot both occupy the same plane. We therefore conjugate the plane located approximately midway between the two mirrors to the objective back focal plane. This distributes the residual conjugation error between the two scanning axes and minimizes pupil displacement during two-dimensional scanning. To determine the correct axial position of the scanning unit, the objective is temporarily removed after carefully marking its position for subsequent reinstallation. A camera (Thorlabs DCC1545M) is then placed at the front focal plane of the tube lens, thereby imaging the objective back focal plane. To ensure the right positioning, the alignment beam is directed through the tube lens (Figure 2c), and the camera is translated along the optical axis until the focused spot reaches its minimum size (see Figure 3b-c). Then, we moved the laser source before the galvanometric mirrors and reduced the beam size with a pinhole (Figure 2d). The two galvanometric mirrors are then driven with sinusoidal voltages of equal effective angular amplitude and with a relative phase shift of 90°. Under these conditions, a correctly conjugated scanner produces a circular beam trajectory in the objective back focal plane, meaning the residual scanning is identical for both mirrors. Instead, if the two mirror pivot planes are unequally conjugated, the trajectory appears elliptical. The complete galvanometric unit is translated along the optical axis until the two axes of the measured trajectory become equal, indicating that the intermediate plane between the mirrors is conjugated to the objective back focal plane (Figure 3e-f). Before applying this criterion, the drive amplitudes should be adjusted to compensate for any difference in the angular response of the two galvanometric axes; otherwise, an elliptical trajectory could arise from unequal scan amplitudes rather than from an axial-conjugation error.

**FIGURE 3.**
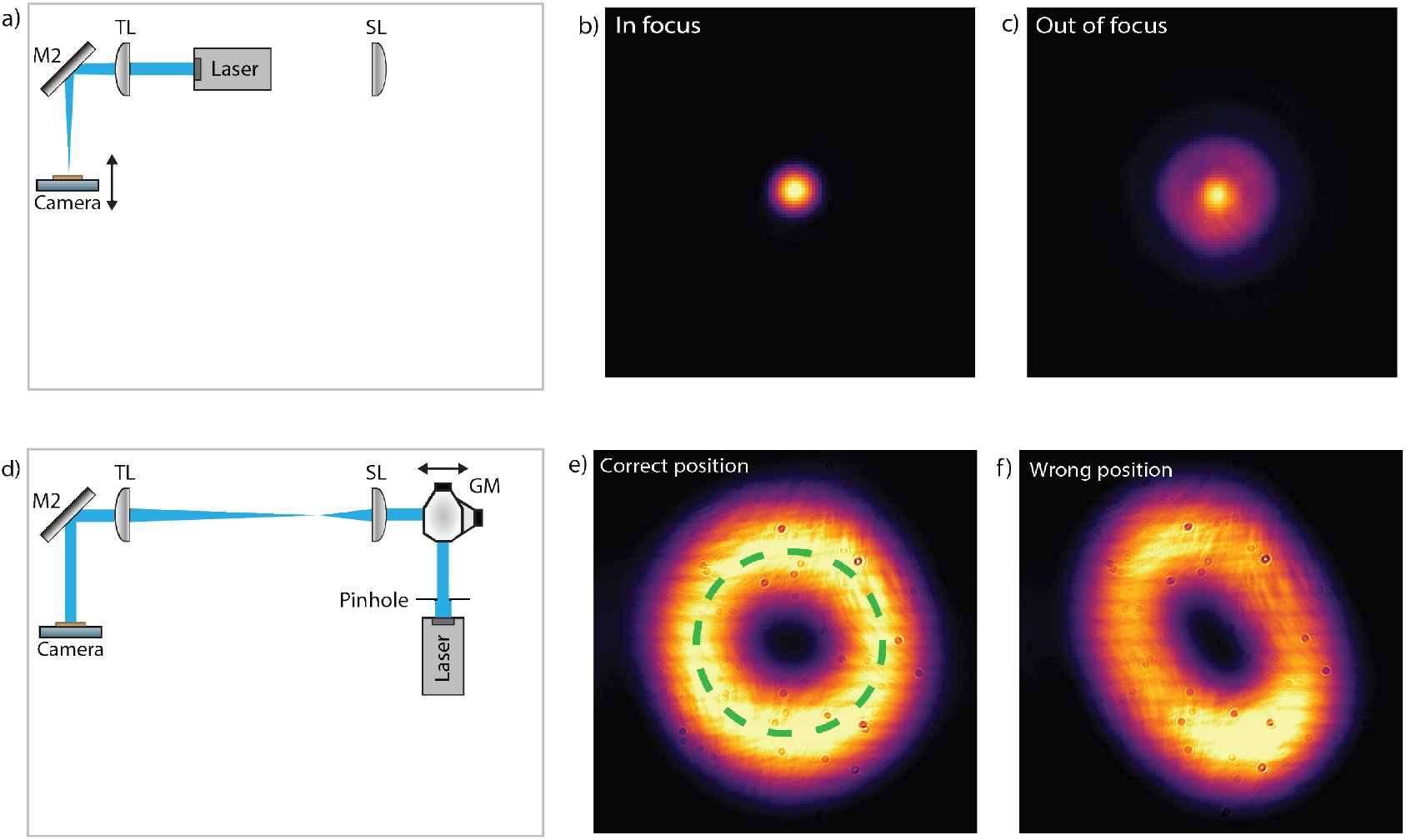
Axial alignment of the galvanometric scanning unit. (a) A camera is positioned at the common focal plane of the objective and tube lens, corresponding to the objective back focal plane after removal of the objective. The camera is translated axially until the alignment-laser spot reaches its minimum size, as illustrated by the in-focus and out-of-focus images in (b) and (c), respectively. (d) The two galvanometric mirrors are driven sinusoidally with equal effective angular amplitudes and a relative phase shift of 90°. The scanning unit is translated along the optical axis while the resulting beam trajectory is observed at the objective back focal plane. A circular trajectory indicates that the plane located approximately midway between the two mirrors is correctly conjugated to the objective back focal plane (e), whereas an elliptical trajectory indicates imperfect conjugation (f).

### 2.2. Aligning the excitation

The dichroic mirror (DM, 25×36 mm Longpass Dichroic Mirror, 490 nm cut-off) is positioned in the microscope assembly. It reflects the 488 nm excitation beam towards the scanning module while trasmitting the fluorescence emitted at longer wavelengths towards the detection path. The excitation source is a fibre-coupled, picosecond-pulsed 488 nm laser module (Flimlabs) connected through a single-mode fibre (Thorlabs P5-405-PCAPC-1). In addition to facilitating beam delivery, the single-modefibre spatiallyfilters the laser output, providing a clean, approximately Gaussian beam profile. The light emerging from the fibre is collimated using an achromatic lens (Thorlabs AC127-019-A, f = 19 mm). The diameter of the collimated beam is determined by the product of the focal length of the collimating lens with the numerical aperture of the fibre. The beam is subsequently expanded by the afocal scan-lens–tube-lens relay according to the ratio between the tube-lens and scan-lens focal lengths. In our configuration, this ratio is 200/50 = 4; therefore, the collimated beam is expanded fourfold before reaching the objective pupil. The collimating lens should be selected by considering both the fibre output divergence and the magnification introduced by the optical relay, such that the beam appropriately fills the objective back aperture. Underfilling the aperture reduces the effective excitation numerical aperture and broadens the focal spot, whereas excessive overfilling causes power loss. If a suitable beam diameter cannot be obtained by changing the collimating lens, an additional beam-expanding telescope can be introduced upstream of the scanning module. Two steering mirrors (M3 and M4) positioned between the laser output and the dichroic mirror are used to direct the excitation beam along the optical axis of the scanning module. Cage-system alignment plates provide an initial reference for centering the beam through the upstream optical components. Final alignment relative to the objective axis is then performed using the two steering mirrors M1 and M2 located between the tube lens and the objective. For this purpose, the objective is temporarily replaced by a tube at least 1 m long, fitted with two diaphragms: one positioned close to the objective mount and the other near the opposite end of the tube. The two spatially separated diaphragms define both the lateral position and the propagation direction of the desired optical axis. M1 and M2 are adjusted by beam walking until the excitation beam passes through the centre of both apertures. This ensures that, when the galvanometric mirrors are in their zero-offset position, the beam is centred on the objective back aperture and propagates perpendicular to its plane. Once this condition is achieved, the alignment tube is removed and the objective is reinstalled at its previously defined position.

### 2.3. Aligning the detection

The detection module consists of the imaging lens (IL; Thorlabs AC254-300-A-ML) with a focal length of 300 mm, a notch filter (Chroma ZET488 Top Notch) for suppressing residual reflected excitation light, and a 7×7 an asynchronous read out SPAD array detector with microlenses (Genoa Instruments). Since the fluorescence is descanned by the same galvanometric mirrors used for excitation, the emitted light forms a stationary beam after the scanning module and can be focused onto the detector array. The imaging lens is positioned relative to the scan lens so that the two lenses form an afocal relay (Figure 2e). The imaging lens then focuses the descanned fluorescence onto the SPAD array. The focal length of this lens determines the additional magnification introduced by the detection relay and must therefore be selected according to the detector size and pixel pitch. In our configuration, the 300-mm imaging lens and the 50-mm scan lens provide a sixfold magnification. Combined with the 100× objective–tube-lens magnification, this results in a total detection magnification of approximately 600×. The detection magnification represents a compromise between spatial sampling and fluorescence collection. It should be sufficiently high so that each SPAD element acts as a sub-Airy-unit pinhole, while allowing a sufficiently large portion of the array to collect most of the fluorescence signal. For an emission wavelength of approximately 500 nm and an objective numerical aperture of 1.4, the Airy-unit diameter in the sample plane is approximately 1.22*λ*/*NA*. At a total magnification of 600×, one Airy unit is mapped onto approximately 261 *µ*m in the detector plane. Because the SPAD-array pitch is 75 *µ*m, each detector element corresponds to approximately 0.29 AU. Consequently, the central 5 × 5 elements span approximately 1.4 AU, whereas the complete 7 × 7 array spans approximately 2 AU. The central 5 × 5 elements were used for the ISM reconstructions presented in this work. This configuration provides sub-Airy-unit sampling at the individual-element level while collectively detecting fluorescence over an area sufficiently large for efficient photon collection.

The image-space depth of focus is approximately given by *DOF*_image_ ≈ 2*nλM* 2/*N A*2, where *n* is the refractive index of the immersion medium, *λ* is the fluorescence wavelength, *M* is the total detection magnification, and *N A* is the objective numerical aperture. For *NA* = 1.4, *n* = 1.5, *M* = 600, and *λ* = 500 nm, the image-space depth of focus is approximately 280 mm. The large detection magnification therefore makes the axial positioning of the detector relatively tolerant. As an initial alignment, the SPAD array can be placed approximately one focal length from the imaging lens. Its final axial position should nevertheless be verified experimentally by comparing the axial positions of the excitation PSF, the central-element detection PSF, and the PSF obtained by summing the detector elements, as described in Section 4.2.

For transverse alignment, the SPAD array is mounted on an XY translation stage (Thorlabs CXYZ1A/M). Its position is adjusted while scanning a thin, signal-rich fluorescent sample and observing the resulting detector fingerprint. The fingerprint is a two-dimensional map of the total signal recorded by each detector element, obtained by integrating the corresponding scanned image over all scan positions. For a sufficiently thin specimen, the fingerprint is theoretically independent of the spatial organization of the labelled structures; the sample therefore does not need to be homogeneous. However, fluorescence should be present over a substantial fraction of the scanned field so that the integrated signal dominates over background and detector dark counts, while out-of-focus fluorescence should be minimized. Suitable samples include a dense layer of fluorescent beads or a cultured cell with an extensively labelled subcellular structure, such as the microtubule network. A centered detector produces an approximately symmetric fingerprint with its photon-weighted centroid close to the central element. The detector position is adjusted until this condition is achieved. Background and detector-specific dark counts, particularly those associated with hot pixels, should be considered when calculating the centroid.

### 2.4. Connection and data collection

Data acquisition in SPAD array-based laser-scanning microscopy must maintain an accurate correspondence between the scan position, detector element, and photon-arrival time while handling the high overall data rate generated by the array. Our microscope is controlled using BrightEyes-MCS [18], an open-source microscope-control and data-acquisition software designed for photon-resolved laser-scanning microscopy with SPAD array detectors. The system supports photon-detection rates in the MHz range while maintaining deterministic synchronization at microsecond pixel dwell times. Here, we summarize the architecture used in the present implementation; further details on its design and operation are provided on the dedicated GitHub pages and in the accompanying publication [18].

The acquisition architecture comprises three principal layers: (i) a National Instruments (NI) FPGA-based general purpose DAC board, (ii) a custom low-level firmware (BrightEyes-MCSLL, https://github.com/VicidominiLab/BrightEyes-MCSLL), and (iii) a high-level control software (BrightEyes-MCS, https://github.com/VicidominiLab/BrightEyes-MCS) which runs on the host computer and provides the user interface, acquisition configuration, data visualization, and storage.

The SPAD array detector is interfaced with the FPGA-based DAC through the PRISM-Light detector-interface unit (Genoa Instruments), which conditions and routes the high-speed digital signals generated by the individual detector elements. This interface allows the FPGA to acquire the detector channels in parallel while preserving their temporal information. The FPGA board (NI PCIe-7820) performs all time-critical operations, including photon counting, scan generation and synchronization, laser-clock handling, and assignment of the detected events to the corresponding spatial and temporal bins. Because these operations are implemented directly in the FPGA hardware, their timing is deterministic and does not depend on the scheduling of the host operating system.

The FPGA is connected to an Analog Front-End (AFE; Genoa Instruments Srl), which provides three analog input channels and five analog output channels. This external interface is required because the NI PCIe-7820 FPGA board, unlike comparable multifunction FPGA boards from National Instruments, does not include integrated analog inputs or outputs. Instead, it provides a larger number of digital I/O lines, which could be used to acquire the 49 parallel digital channels generated by the SPAD array. The AFE complements these digital capabilities by providing the analog outputs required to control the galvanometric mirrors and piezoelectric stage, as well as analog inputs for auxiliary signals such as photomultiplier-tube outputs and scanner-position feedback.

The BrightEyes-MCSLL firmware, implemented in LabVIEW FPGA, uses parallel photon-counting pipelines and buffered data transfer through direct-memory-access first-in,first-out buffers (DMA FIFOs). This architecture supports continuous acquisition without observable data loss, provided that the detector count rate and aggregate data throughput remain within the validated operating limits of the system. The scan-control and synchronization signals are generated at the FPGA level, ensuring precise spatiotemporal correspondence among modulated or pulsed excitation, scanning, and detection.

The acquired data are stored in HDF5 format together with the acquisition metadata and FPGA configuration. This format preserves the multidimensional structure of the ISM dataset, including the scan coordinates, detector-element index, and, when enabled, photon-arrival-time bin.

### 3. MICROSCOPE CALIBRATION

After optical assembly and alignment, the microscope must be calibrated to relate the electronic control signals to physical displacements in the sample. We therefore determine the voltage-to-displacement conversion factors of the galvanometric scanners and the piezoelectric Z actuator, establishing the spatial sampling along all three dimensions. These calibrations are essential for defining the image pixel size,field of view, and axial spacing of three-dimensional image stacks.

### 3.1. Calibrating the galvonometers translation scale

The galvanometric mirrors are controlled by analog voltages generated by the digital-to-analog converters of the Genoa Instruments analog front-end (AFE). The corresponding digital control values are calculated by the BrightEyes-MCSfirmware and transmitted by the FPGA board to the AFE, which converts them into the analog voltages applied to the galvanometer drivers. These voltages determine the angular deflections of the mirrors, which are converted by the scan-lens–tube-lens–objective relay into lateral displacements of the excitation focus in the sample plane. The resulting voltage-to-displacement relationship depends on the galvanometer response, the optical-relay magnification, and the system alignment, and must therefore be determined experimentally. This calibration is required to assign physical dimensions to the acquired images, define the scan pixel size and field of view.

We determine the conversion factors using the three-dimensional fluorescent ring matrix of an Argo-SIM v1 calibration slide (Argolight), in which the ring centers are separated by a known distance of 5 *µ*m. The fluorescent reference structure must be compatible with the excitation and detection wavelengths used during calibration. The slide employed here is designed for 488 nm excitation; however, the resulting voltage-to-displacement calibration can also be applied to other excitation wavelengths available on the same microscope, because this relationship is primarily determined by the galvanometer response and the geometry of the scanning relay rather than by wavelength. This remains valid provided that the same objective and scanning optics are used and that wavelength-dependent changes in the beam path or relay magnification are negligible.

An XY image of a single ring layer is acquired over a voltage range representative of routine imaging (Figure 4a). A relatively long pixel dwell time and moderate scan speed are used to minimize positional lag and settling errors. The calibration structure is positioned near the centre of the field of view, away from the acceleration, deceleration, and turnaround regions at the scan boundaries. The rings are segmented using unsupervised clustering, and their centers are fitted to a rectangular grid with the known physical spacing (Figure 4b). The fit provides independent conversion factors, in micrometres per volt, for the X and Y axes. Using a single conversion factor for each axis assumes a linear voltage-to-displacement relationship, which is generally valid for small deflection angles within the central scanning range. The fit residuals should nevertheless be inspected to verify this assumption and to identify possible axis non-orthogonality or field-dependent distortions. If appreciable deviations from linearity are observed, the calibration should be repeated over a smaller, more central field of view.

**FIGURE 4.**
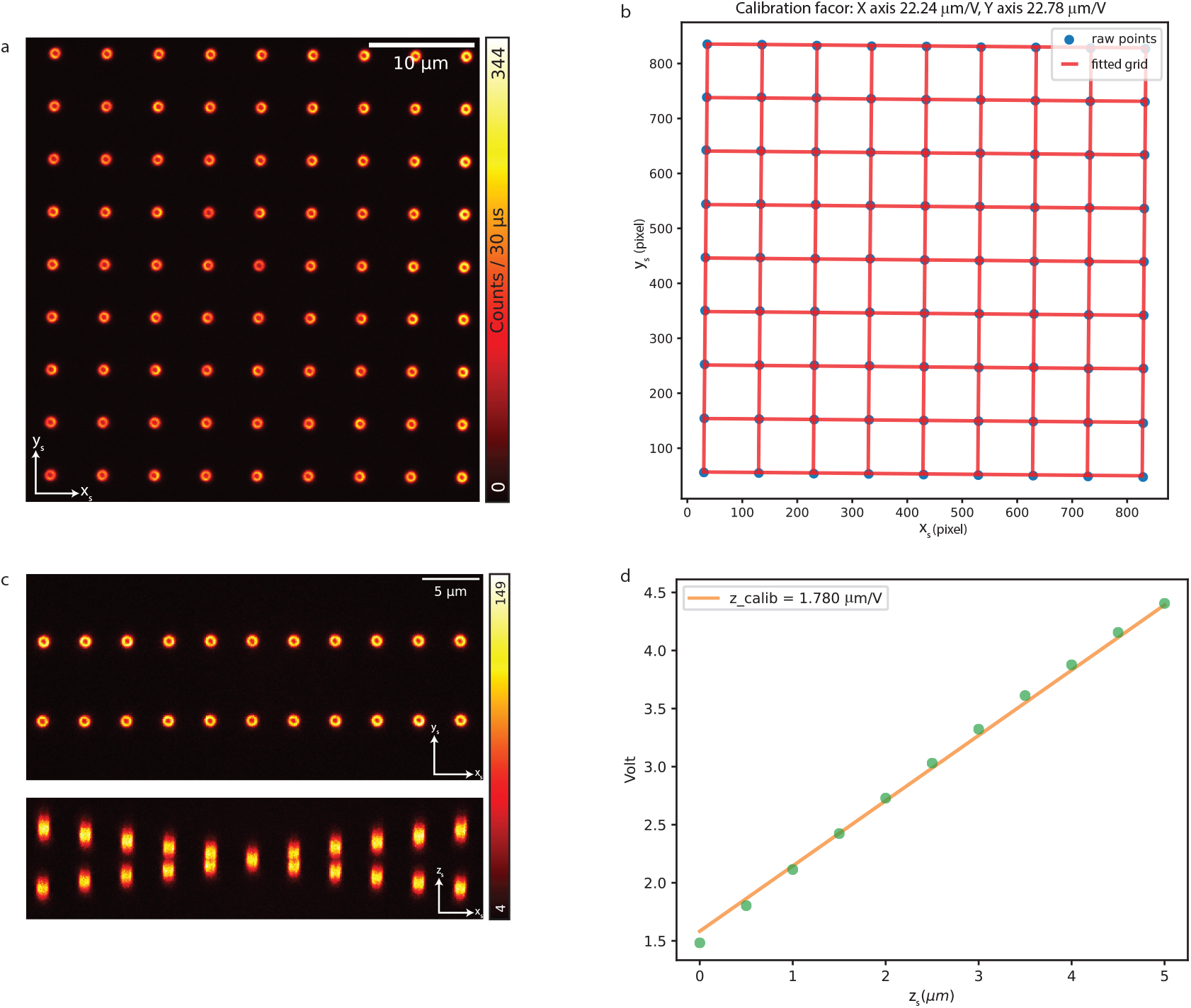
Calibrations of XYZ displacement. a-b: Calibration of the galvanometer mirrors voltage for XY scanning of the laser beam. a: Sum fluorescence image of rings matrix spaced by 50 *µm*. b: After localization of each rings, the matrix is fitted by a rectangular grid with two different pitches in x and y directions to calculate the calibration factors of the galvanometric mirrors. c-d: Calibration of the piezoelectric stage for Z displacement of the sample. c: Sum fluorescence image of rings in stairs arrangement with 500 nm step. d: Voltage applied on the piezoelectric stage driver against the axial localization of the rings

### 3.2. Calibrating the piezoelectric stage axial displacement

The fine axial position of the sample is controlled by the piezoelectric Z actuator of the three-axis NanoMax flexure stage. The FPGA generates the digital control signal, which is converted into an analog voltage by the Genoa Instruments AFE and delivered to the piezoelectric-stage controller (Thorlabs MDT693B). The relationship between the applied voltage and the resulting sample displacement must be calibrated experimentally to assign a physical axial spacing to acquired image stacks and to measure three-dimensional PSFs and sample structures accurately.

We determine the axial conversion factor using the fluorescent 3D crossing-stair structure of the Argo-SIM v1 calibration slide (Argolight), which contains steps separated axially by 500 nm. A 3D image stack is acquired over a voltage range representative of routine imaging (Figure 4c). The fluorescent stair structures are segmented into individual steps using unsupervised clustering, and the axial center of each step is estimated from its fluorescence distribution. The measured axial positions are then fitted to a linear model using the known 500 nm spacing, providing the Z-actuator conversion factor in micrometres per volt (Figure 4d). The two crossing stair structures are analysed independently, providing an internal consistency check of the calibration.

The calibration should be performed over a voltage range and under acquisition conditions representative of routine imaging. The residuals of the linear fit should be inspected to verify that a single conversion factor adequately describes the selected operating range. This procedure can be readily adapted to other voltage-controlled Z-axis actuators.

## 4. MICROSCOPE CHARACTERIZATION

Following spatial calibration, the optical performance of the microscope must be experimentally characterized to verify that the assembled system operates as intended. We first measure the detection magnification, which determines the detector sampling in Airy units and is required for quantitative ISM reconstruction. We then characterize the excitation and detection PSFs to assess image quality and verify the spatial overlap and alignment of the excitation and detection paths.

### 4.1. Magnification characterization

A defining feature of an ISM system is the high detection magnification, denoted by *M*, required to map each detector element onto a sub-Airy-unit region in the sample plane while allowing the array to cover the desired total detection area. Measuring the actual magnification is therefore important for determining the detector sampling in Airy units, verifying the optical design, and providing an accurate spatial model for ISM reconstruction.

To measure *M*, the microscope is converted into a wide-field-like configuration by replacing the SPAD array with a camera (Thorlabs DCC1545M) positioned in the same image plane (Figure 5a). In principle, the SPAD array could be used directly because it is also a spatially resolved detector. However, its small number of elements and limited field of view provide insufficient sampling for a robust magnification measurement. The dichroic mirror is mounted in a 30 mm cage cube designed for fluorescence filter sets, allowing it to be removed and reinserted rapidly while preserving repeatable alignment. For this measurement, it is temporarily removed to maximize the transmitted-light intensity reaching the camera and subsequently reinserted without requiring realignment of the microscope.

**FIGURE 5.**
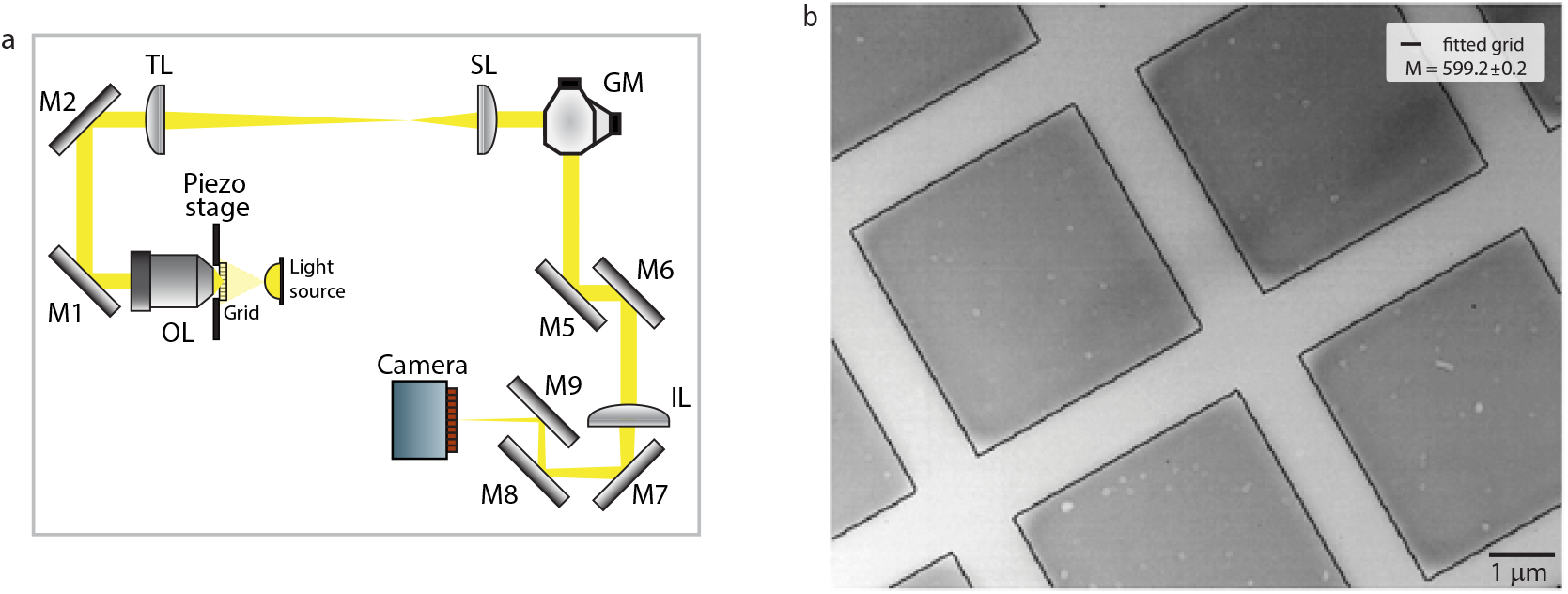
Measurement of the detection magnification. (a) Schematic of the wide-field-like configuration used for the measurement. The SPAD array is replaced by a camera positioned in the same image plane. The dichroic mirror, mounted in a 30 mm cage cube with an interchangeable filter-set holder, is temporarily removed to maximize the transmitted-light intensity reaching the camera while enabling repeatable repositioning. (b) Transmission image of the calibration grid with the fitted grid model overlaid.

A calibration grid with a known pitch of 5 *µ*m (Thorlabs R1L3S3P1) is placed in the sample plane and illuminated in transmission using a lamp. Since the expected magnification is approximately 600×, a small grid pitch is required to ensure that multiple periods remain visible within the camera field of view. The resulting grid image is fitted with a rectangular-grid model (Figure 5b). From the fitted grid spacing in camera pixels and the known camera pixel size of 5.20 *µ*m, we determine the grid period in the detector plane. The ratio between this value and the known 5 *µ*m period in the sample plane yields the detection magnification. Using this procedure, we measure *M* = 599.2 ± 0.2 (mean ± standard error from 100 acquired images), in close agreement with the nominal magnification of 600×. The measurement also confirms that the detection relay maps one Airy unit and the SPAD-array pitch onto the expected dimensions in the detector and sample planes, respectively.

### 4.2. Point-spread functions characterization

Following transverse alignment of the SPAD array, the excitation and effective detection PSFs are measured to assess image quality and verify their three-dimensional registration. For completeness, Figure 6a–b shows representative fingerprints acquired before and after the transverse-alignment procedure described in Section 2.3. The remaining panels summarize the measurements used to characterize the excitation and detection PSFs.

**FIGURE 6.**
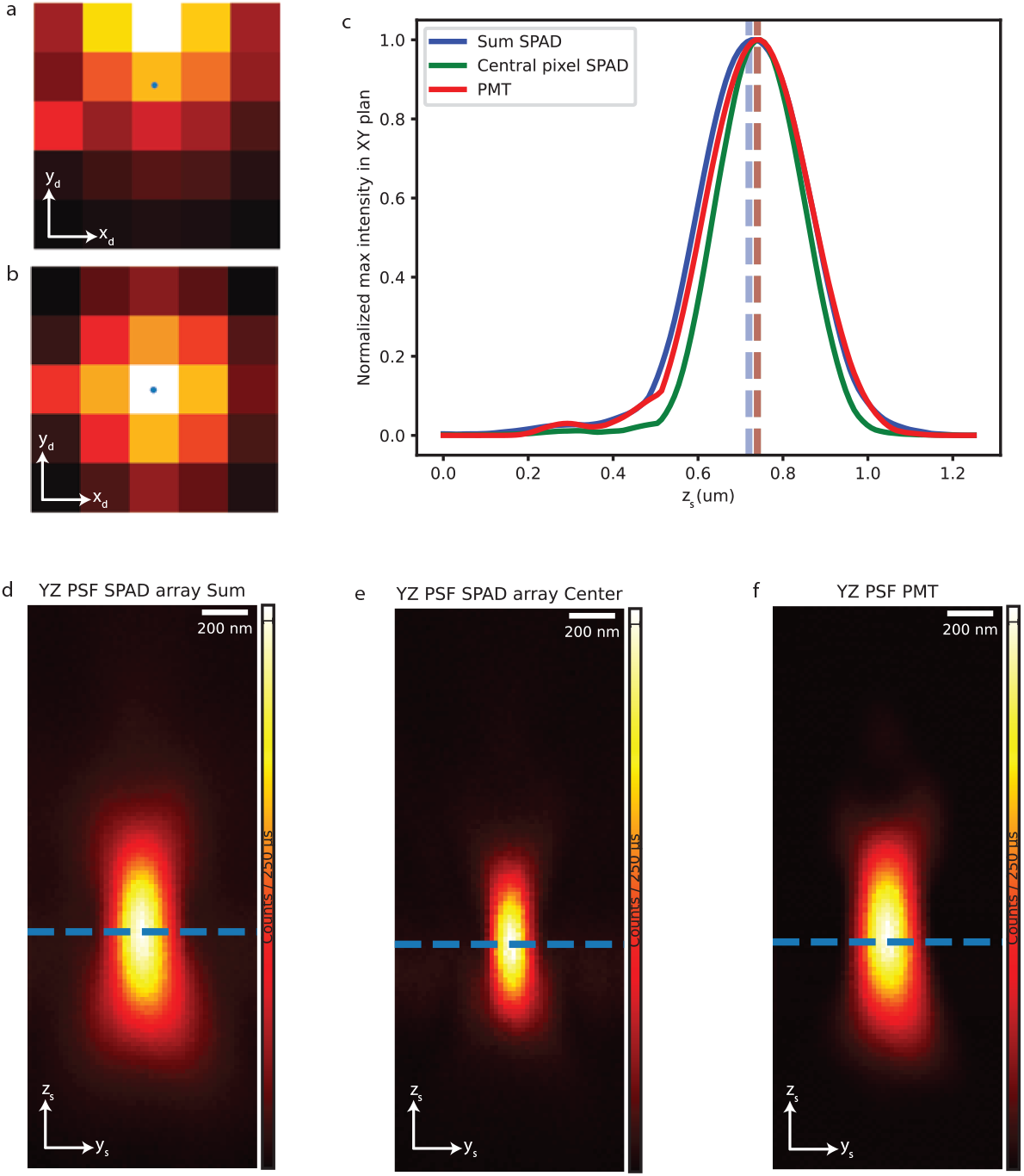
Detector alignment and PSF characterization using scattered light. (a,b) Detector fingerprints obtained by integrating the signal recorded by each SPAD element over the complete scan. The blue marker indicates the photon-weighted centroid of thefingerprint. A displaced centroid reveals transverse misalignment of the detector (a), whereas a centroid located near the central element indicates correct alignment (b). The element immediately to the left of the central pixel exhibits an abnormally high count rate. (c–e) YZ projections of a 3D scan of a 100-nm gold nanoparticle: signal summed over all SPAD-array elements (c), signal recorded by the central SPAD element (d), and non-descanned PMT signal (e). (f) Normalized maximum intensity in each XY plane as a function of the axial sample position for the three detection configurations.

In an ISM system, the image recorded by each detector element is governed by the product of the excitation PSF and the spatially shifted detection PSF associated with that element. It is therefore important to evaluate the excitation PSF, the effective PSF measured by the SPAD array, and their spatial registration.

The excitation PSF is characterized using 100-nm gold nanoparticles and non-descanned detection. A three-dimensional XYZ scan of an isolated nanoparticle is acquired in scattering mode. A pellicle beam-splitter (Thorlabs BP145B1) inserted between the objective and tube lens directs part of the scattered light towards a photomultiplier tube (PMT). Because the PMT collects the scattered light without spatial pinholing, its signal primarily reflects the spatial distribution of the excitation intensity, convolved with the finite size of the nanoparticle. The resulting 3D distribution therefore provides an experimental approximation of the excitation PSF. An asymmetric or distorted PSF can indicate imperfect excitation alignment or optical aberrations.

The light transmitted through the pellicle is simultaneously recorded by the SPAD array, providing the effective PSF of the ISM detection path. The YZ or XZ projections of the signal summed over the array, the central-element signal, and the non-descanned PMT signal are compared (Figure 6c–f). Coincidence of their axial maxima confirms that the excitation focus, the central-element detection PSF, and the effective array-detection plane are correctly registered. An axial displacement between these maxima indicates that the position of the SPAD array or imaging lens should be adjusted.

Finally, the detection PSF can be inspected directly from the SPAD-array micro-image. The excitation beam is centred on an isolated gold nanoparticle and the scan amplitude is set to zero, so that the signal distribution across the detector elements provides a sparsely sampled image of the detection PSF. Its position, symmetry, and extent offer an additional check of detector centring and of the detection magnification.

## 5. IMAGE ACQUISITION AND DATA ANALYSIS

After calibration and optical characterization, the acquisition parameters must be selected to preserve both the spatial information encoded by the scan and the linear photon-counting response of the detection system. The raw ISM dataset consists of one scanned image for each detector element and therefore contains both the sample-space co-ordinates defined by the excitation scan and the detector-space coordinates provided by the SPAD array.

The scan pixel size of the raw dataset should be selected according to the highest spatial frequency that the reconstructed ISM image is expected to contain. Similar to an ideal confocal microscope with an infinitesimally small pinhole, ISM has an effective PSF given by the product of the excitation and detection PSFs. Its optical transfer function therefore has a cut-off frequency approximately equal to the sum of the excitation and detection cut-off frequencies. For similar excitation and emission wavelengths, this cut-off frequency is approximately twice that of conventional wide-field imaging. In practice, an ideal confocal microscope cannot efficiently exploit this extended bandwidth because the extremely small pinhole required to reach it rejects most of the fluorescence and markedly reduces the signal-to-noise ratio. ISM overcomes this limitation by recording the sub-Airy-unit signal separately on each detector element and subsequently combining the detected photons computationally. It can therefore approach the cut-off frequency of an ideal confocal microscope while maintaining high photon-collection efficiency. The spatial sampling of the raw scan should consequently support this extended frequency range to prevent aliasing during reconstruction. More generally, the ISM cut-off frequency can be approximated as *f*_c,ISM_ ≈ 2*NA* (1/*λ*exc + 1/*λ*em). According to the Nyquist criterion, the scan pixel size should therefore satisfy Δ*x*scan ≤ 1/(2*f*_c,ISM_). When the excitation and emission wavelengths are approximated by a common wavelength *λ*, this condition reduces to Δ*x*scan ≲ *λ*/(8*N A*). For example, with 488-nm excitation, fluorescence emission around 520 nm, and an objective with *N A* = 1.4, a scan pixel size of approximately 50 nm or smaller is appropriate. This represents a conservative acquisition criterion that prevents the scan sampling from limiting the spatial frequencies available to the reconstruction. Some ISM algorithms can accommodate larger scan pixels by exploiting the complementary subpixel shifts among the detector-element images and resampling the reconstructed image on a finer grid [9]. This procedure uses the additional spatial information encoded by the detector array and should not be confused with simple numerical interpolation, which cannot recover spatial frequencies lost through undersampling. For the reference implementation described here, a scan pixel size of approximately 50 nm or smaller remains the most generally applicable choice.

Detector saturation must be avoided irrespective of the reconstruction algorithm. At high photon fluxes, the dead time of each SPAD produces a nonlinear relationship between the incident photon flux and the measured count rate, distorting the individual detector images. This violates the linear image-formation model assumed by model-based reconstruction methods and can also introduce artefacts into approaches such as pixel reassignment, even though they do not explicitly invert this model. Since the central detector element generally receives the highest photon flux, its count rate provides a conservative indicator of detector saturation. For the SPAD array used here, each element has a dead time of approximately 30 ns, corresponding to a theoretical maximum count rate of about 33 Mcps. Operation well below this limit is nevertheless required to maintain an approximately linear response. As a practical criterion, the count rate of the central element should preferably remain below approximately 1 Mcps, corresponding to a dead-time occupancy of about 3%.

Detector saturation must be distinguished from saturation of the data-transfer pipeline. Distributing the fluorescence over multiple detector elements reduces the count rate per element and therefore mitigates SPAD saturation, but simultaneously increases the total data rate that must be transferred and stored. BrightEyes-MCS monitors this condition and reports, for each scan pixel, an overflow alarm when the data-transfer capacity is exceeded. Acquisitions containing such alarms should not be used for quantitative analysis or image reconstruction. The excitation power, pixel dwell time, and acquisition settings should therefore be adjusted until both the central-element count rate remains within the selected linearity range and no data-transfer overflow is reported.

Several algorithms are available for reconstructing a super-resolved image from an ISM dataset [6, 19, 20, 7, 8, 11]. For an array comprising *N* active detector elements, the dataset consists of *N* scanned images, each associated with a different detector position. In our implementation, the central 5 × 5 elements of the SPAD array are used, resulting in 25 scanned images (Figure 7a). The most widely used reconstruction approaches are based on pixel reassignment [19]. Each detector element records a similar image of the specimen, laterally displaced according to its position within the array and differing from the other images primarily in signal level and noise. Pixel reassignment consists of estimating these detector-dependent displacement vectors, shifting each image in the opposite direction, and summing the registered images. In adaptive pixel reassignment (APR) [7], the displacement vectors are estimated directly from the acquired dataset by calculating the phase correlation between each detector image and a reference image, typically the image recorded by the central element. The resulting shift vectors are shown in Figure 7b.

**FIGURE 7.**
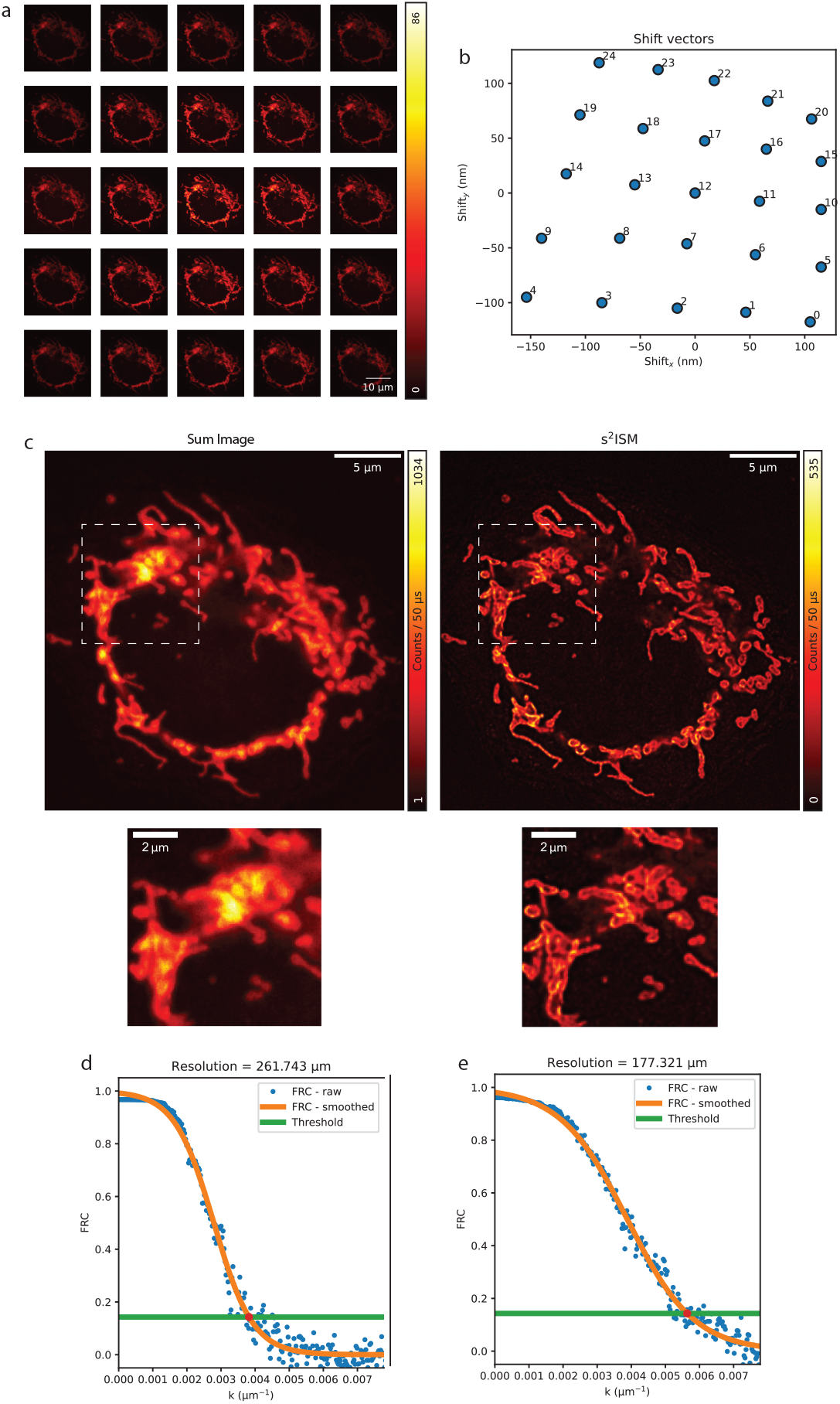
Image processing. HeLafixed (PFA 4% in Prolong Diamond) cell with TOMM20 labelled with AF488. a: Scanned images. b: Calculated shift vectors with cross-correlations between the scanned images. c: Sum image of the 25 scanned images, considered as a confocal open pinhole (1.4 airy units) equivalent image, compared to the in-focus s^2^ISM image. d: Fourier Ring Correlation (FRC) computation of the confocal open pinhole image. e: FRC of the in-focus s^2^ISM image.

As an alternative to pixel-reassignment methods, we recently introduced super-resolution optical-sectioning ISM (s^2^ISM) [11], a model-based reconstruction algorithm that inverts the physical image-formation model of ISM (https://github.com/VicidominiLab/s2ISM). The method provides the lateral-resolution enhancement achieved by conventional ISM reconstruction while additionally separating the in-focus signal from the out-of-focus background, thereby restoring optical sectioning. In Figure 7c, the in-focus s^2^ISM reconstruction of a fixed HeLa cell immunolabelled for TOMM20 reveals the hollow structure of individual mitochondria. Since TOMM20 localizes to the mitochondrial outer membrane, this appearance becomes discernible through the combined improvement in lateral resolution and optical sectioning. The Fourier ring correlation (FRC) in Figure 6d–e shows a resolution improvement of 1.47. From theory, we should expect a two-fold enhancement of cut-off frequency. However this ideal case without noise is when we compare the super-resolved image to the no-pinhole detection one, whereas here the sum image is already a confocal system with enhanced resolution with a pinhole of approximately 1.2 AU.

## 6. DISCUSSION AND CONCLUSION

We presented BrightEyes-Scope, a reference implementation of ISM based on SPAD-array detection. Its optical architecture remains closely related to that of a conventional confocal laser-scanning microscope, facilitating both construction from standard optical components and the conversion of existing custom-built systems. The principal differences lie in the replacement of the conventional pinhole and single-element detector with a SPAD array and in the detection magnification required to map each detector element onto a sub-Airy-unit region of the sample. In our implementation, the relatively high detection magnification is determined by the 75*µ*m pixel pitch of the array. Each SPAD element acts as an independent confocal detection channel, while the complete array collects fluorescence over a larger detection area. Computational reconstruction combines these signals to achieve the resolution expected from an ideal confocal laser-scanning microscope – that is, up to a twofold improvement in lateral resolution over diffraction-limited wide-field microscopy – without incurring the severe photon loss associated with an almost-closed confocal pinhole. The implementation described here emphasizes the practical aspects that determine ISM performance, including detection magnification, detector centring, registration of the excitation and detection PSFs, and scan sampling. Proper optimization of these parameters ensure that the acquired dataset contains the spatial information required to achieve the expected twofold improvement in lateral resolution after ISM reconstruction. The procedures presented here are therefore intended not only to support the construction of BrightEyes-Scope but also to provide a general framework for diagnosing and optimizing other array-based ISM implementations.

The microscope presented here uses 488-nm excitation, detection in the green spectral range, a high-numerical-aperture oil-immersion objective, and a 7 × 7-element SPAD array from Genoa Instruments. These choices represent a specific implementation rather than intrinsic limitations of the architecture. The same design principles can be adapted to other wavelengths, objectives, scanning configurations, and asynchronous SPAD arrays, including the 23-element SPAD23 from Pi Imaging, whose elements are arranged on a hexagonal lattice with a 23 *µ*m nearest-neighbour pitch. Its smaller pitch permits a proportionally lower detection magnification while preserving the same sub-Airy-unit sampling. The modular architecture also permits the incorporation of additional excitation sources and detection channels. BrightEyes-Scope is controlled through the open-source BrightEyes-MCS hardware and software microscope control platform [18]. Together with the shared optical design, calibration procedures, reconstruction software, and example datasets, it provides a reproducible starting point for laboratories wishing to construct an ISM system or upgrade an existing laser-scanning microscope. We anticipate that this reference implementation will lower the technical barrier to SPAD-array-based ISM and facilitate its adoption as a photon-efficient, super-resolution extension of confocal microscopy.

The capabilities of BrightEyes-Scope combined with BrightEyes-MCS extend beyond ISM. BrightEyes-MCS implements multichannel time-to-digital conversion based on the Digital Frequency Domain (DFD) approach [10] on the same National Instruments FPGA board used to control the microscope, thereby enabling fluorescence-lifetime imaging microscopy (FLIM). This implementation bins detected photons according to their arrival time relative to the excitation laser and generates photon-arrival-time histograms directly during acquisition. The FLIM-enabled version of BrightEyes-MCS will be described in a subsequent publication. We also plan to extend the BrightEyes-Scope and BrightEyes-MCS to fluorescence fluctuation spectroscopy [16] and, in the longer term, to real-time single-particle tracking [21] and scanning-based single-molecule localization microscopy [22, 23].

A further development will be the integration of BrightEyes-MCS with the BrightEyes-TTM multichannel time-tagging module [14]. Whereas the FPGA-based DFD implementation reduces the data during acquisition by binning and histogramming photon-arrival times, BrightEyes-TTM records individual photon time tags and therefore preserves the complete temporal information associated with each detected event. Combining these complementary acquisition modes within a unified framework will enable users to select between efficient real-time processing and fully time-tagged acquisition according to the requirements of the experiment.

BrightEyes-Scope provides the open and reproducible optical-hardware foundation of a broader open-science project aimed at making photon-resolved microscopy accessible to and widely adopted by the scientific community.

## Acknowledgements

We thank G. Garrè (Istituto Italiano di Tecnologia, Genoa, Italy) for providing useful guidance for data analysis. We thank the French Research Group Imabio for giving us the opportunity to present the microscope architecture of BrightEyes-Scope during the thematic school MiFoBio 2025.

## Supporting Information

The codes for calibration are available as a Python notebook at https://github.com/VicidominiLab/BrightEyes-Scope.git. The algorithm for data analysis with s^2^ISM algorithm is available at https://github.com/VicidominiLab/s2ISM/tree/main. All the raw data and 3D CAD of the setup (rendered with Fusion 360) are available at https://doi.org/10.5281/zenodo.20204690.

## references

[1] Lichtman JW, Conchello JA. Fluorescence microscopy. Nat Methods 2005;2(12):910–919.

[2] Neil MAA, Juškaitis R, Wilson T. Real time 3D fluorescence microscopy by two beam interference illumination. Optics Communications 1998;153(1):1–4.

[3] Gustafsson M. Surpassing the lateral resolution limit by a factor of two using structured illumination microscopy. Journal of Microscopy 2000;198(2):82–87.

[4] Wilson T. Optical sectioning in confocal fluorescent microscopes. J Microsc (Oxford, U K) 1989;(154):143–156.

[5] Sheppard CJR, Cogswell CJ. Three-dimensional image formation in confocal microscopy. J Microsc (Oxford, U K) 1990;159(2):179–194.

[6] Müller CB, Enderlein J. Image Scanning Microscopy. Physical Review Letters 2010;104(19):198101.

[7] Castello M, Tortarolo G, Buttafava M, Deguchi T, Villa F, Koho S, et al. A robust and versatile platform for image scanning microscopy enabling super-resolution FLIM. Nature Methods 2019;16(2):175–178.

[8] Zunino A, Castello M, Vicidomini G. Reconstructing the image scanning microscopy dataset: an inverse problem. Inverse Problems 2023;39(6):22.

[9] Zunino A, Slenders E, Fersini F, Bucci A, Donato M, Vicidomini G. Open-source tools enable accessible and advanced image scanning microscopy data analysis. Nature Photonics 2023;17(6):457–458.

[10] Tortarolo G, Zunino A, Piazza S, Donato M, Zappone S, Pierzyńska-Mach A, et al. Compact and effective photon-resolved image scanning microscope. Advanced Photonics 2024;6(1):016003.

[11] Zunino A, Garrè G, Perego E, Zappone S, Donato M, Vastenhouw N, et al. Structured detection for simultaneous super-resolution and optical sectioning in laser scanning microscopy. Nature Photonics 2025;(8):888–897.

[12] Huff J. The Airyscan detector from ZEISS: confocal imaging with improved signal-to-noise ratio and super-resolution. Nat Methods 2015;12(12):1–2.

[13] Buttafava M, Villa F, Castello M, Tortarolo G, Conca E, Sanzaro M, et al. SPAD-based asynchronous-readout array detectors for image-scanning microscopy. Optica 2020;7(7).

[14] Rossetta A, Slenders E, Donato M, Zappone S, Fersini F, Bruno M, et al. The BrightEyes-TTM as an open-source time-tagging module for democratising single-photon microscopy. Nature Communications 2022;(1):7406.

[15] Slenders E, Perego E, Buttafava M, Tortarolo G, Conca E, Zappone S, et al. Cooled SPAD array detector for low light-dose fluorescence laser scanning microscopy. Biophys Rep 2021;1(2):1–13.

[16] Slenders E, Castello M, Buttafava M, Villa F, Tosi A, Lanzanò L, et al. Confocal-based fluorescence fluctuation spectroscopy with a SPAD array detector. Light: Science & Applications 2021;10(1):31.

[17] Perego E, Zappone S, Castagnetti F, Mariani D, Vitiello E, Rupert J, et al. Single-photon microscopy to study biomolecular condensates. Nat Commun 2023;14(1):8224.

[18] Donato M, Slenders E, Zunino A, Bega L, Vicidomini G. BrightEyes-MCS: a control software for multichannel scanning microscopy. Advanced Photonics 2024;9(103):7125.

[19] Sheppard CJR, Shalin BM, Heintzmann R. Superresolution by image scanning microscopy using pixel reassignment. Optics Letters 2013;38:2889–2892.

[20] Gregor I, Enderlein J. Image scanning microscopy. Current Opinion in Chemical Biology 2019;51:74–83.

[21] Bucci A, Tortarolo G, Held MO, Bega L, Perego E, Castagnetti F, et al. 4D Single-particle tracking with asynchronous read-out single-photon avalanche diode array detector. Nature Communications 2024;15(1):6188.

[22] Radmacher N, Nevskyi O, Gallea JI, Thiele JC, Gregor I, Rizzoli SO, et al. Doubling the resolution of fluorescence-lifetime single-molecule localization microscopy with image scanning microscopy. Nat Photon 2024;18(10):1059–1066.

[23] Slenders E, Patil S, Held MO, Zunino A, Vicidomini G. Array detection enables large localization range for simple and robust MINFLUX. Light: Science & Applications 2025;14(1):234.

